# Temporal and spatial localization of prediction-error signals in the visual brain

**DOI:** 10.1101/079848

**Authors:** Patrick Johnston, Jonathan Robinson, Athanasios Kokkinakis, Samuel Ridgeway, Michael Simpson, Sam Johnson, Andrew W. Young

**Affiliations:** Institute of Health and Biomedical Innovation, Queensland University of Technology, Australia; York Neuroimaging Centre and Department of Psychology, University of York, UK

**Keywords:** Predictive coding, visual perception, EEG, MEG, N170

## Abstract

It has been suggested that the brain pre-empts changes in the visual environment through generating predictions, although real-time eletrophysiological evidence of prediction violations remains elusive. In a series of experiments we showed participants sequences of images that followed a predictable implied sequence or whose final image violated the implied sequence. Through careful design we were able to use the same final image transitions across predictable and unpredictable conditions, ensuring that any differences in neural responses were due only to preceding context and not to the images themselves. EEG and MEG recordings showed that early/mid-latency visual evoked potentials were robustly modulated by images that violated the implied sequence across a range of types of image change (expression deformations, rigid-rotations and visual field location). This modulation occurred irrespective of stimulus object category. Although the stimuli were static images, MEG source reconstruction of the early latency signal (N/M170) localised expectancy violation signals to brain areas associated with motion perception. Our findings suggest that the N/M170 can index mismatches between predicted and actual visual inputs in a system that predicts trajectories based on ongoing context. This has important implications for understanding the N/M170 and investigating how the brain represents context to generate perceptual predictions.

## Introduction

It has long been recognized that top-down influences play a role in perception. An influential refinement of this idea is that rather than passively registering sensory data, the brain is hypothesized to actively generate and test predictions about its likely sensory input on a moment-by-moment basis (Gregory, 1980). Models of perceptual prediction therefore focus upon the need for mechanisms that attempt to minimize prediction error within reciprocally interconnected hierarchical networks (Friston & Kiebel, 2009; Panichello et al, 2012; Summerfield & De Lange, 2014). Behaviorally, there is growing support for the existence of such mechanisms. For instance, the phenomenon of *representational momentum* suggests the existence of dynamically evolving representations that model object trajectories (Hubbard, 2005), including biological motion trajectories (Kaufman & Johnston, 2014). Such findings suggest the possibility of identifying brain activity indices reflecting error-checking mechanisms at early stages of visual perception. We propose that the evoked brain response known as the N/M170 may provide such an index.

First reported by Bentin et al. (1996), the N170 Event Related Potential (ERP) has proved to be a robust and highly replicable index of early visual cognition (Johnston et al., 2015). Recorded at occipito-temporal electrodes, the N170 is a negative inflection of the ERP occurring ~150-200ms following stimulus onset. There has been much focus on the N170 (M170 in magnetoencephalography/MEG) (Halgren et al., 2000)) as an index of face-sensitive processes, since the N/M170 is generally larger to faces than to other object categories (Liu et al., 2000; Rossion and Jacques, 2008; Eimer, 2011). However, the N/M170 is also robustly elicited by non-face stimuli including objects of expertise (Tanaka and Curran, 2001) visual word-forms (McCandliss et al., 2003), naked bodies (Hietanen and Nummenmaa, 2011), and conditioned danger signals (Levita et al., 2015).

A common assumption has been that the N/M170 predominantly reflects stimulus-driven processes – indeed the dominant view is that the “face N/M170” indexes the structural encoding of faces preceding facial identification (Eimer, 2011). However, the N/M170 may be subject to influences of top-down modulation (Righart & de Gelder, 2006; Furl et al., 2007; Hietanen & Astikainen, 2013) and visual salience (Hietanen & Nummenmaa, 2011; Levita et al., 2015).

Importantly, the N/M170 is believed to be the first component of the ERP capable of indexing higher-level vision, since earlier components (eg. P1) are not sensitive to stimulus category (Rossion & Caharel, 2011). This makes it a natural potential candidate for indexing predictive mechanisms since the early stages of higher-level vision are likely to involve the resolution of incoming sensory data with top-down influences on perception. However, the standard visual ERP paradigm involves stimuli being selected at random from a fixed set and presented following a near-blank “fixation” screen. Whilst this has advantages in terms of experimental control, it means that every trial is, in effect, a quasi-independent context-free event. By contrast, in everyday life, many aspects of the visual environment are predictable and discontinuities in our visual input are mostly due to natural external properties such as occlusion or internally generated events such as blinks and saccades. Thus, adherence to the standard visual ERP paradigm may have masked some important aspects of the N/M170. We propose that the N/M170 may, in part, index “visual surprise” to the unpredicted appearance of a potentially important stimulus change.

We tested this idea in a series of three EEG experiments and a fourth experiment using MEG source reconstruction. In each experiment participants viewed a sequence of five successive static images on each trial. The first four images in each sequence created a contextual trajectory of implied movement, such as a regular series of changes in position. The final fifth image in each sequence either conformed to, or violated, the expected trajectory. Experiment 1 used facial expression trajectories, Experiment 2 used rigid-body rotation trajectories (of heads and body images) and Experiment 3 used locational trajectories (for faces and shapes). Experiment 4 was identical to the second experiment but was performed in the MEG scanner. Each experiment was conducted using a new sample of participants. The design of each experiment was such that exactly the same pairs of images were used to create predictable or unpredictable final stimulus transitions, to provide compelling demonstrations that any differences in neural responses were directly due to predictability and not to visual properties of the stimuli themselves.

## Methods

### Experiment 1

We first examined facial expression trajectories. Trials consisted of the presentation of sequences of five static images that followed a consistent direction of change in expression across the first four images, and where the final image either did or did not conform to the established pattern. We predicted that if the N170 can serve as an index of predictive mechanisms, amplitudes to Unpredictable final images would be larger than to Predictable final images.

### Participants

There were 20 participants (10 female; mean age 23.5 years, SD 4.0). All were an opportunity sample from the undergraduate and postgraduate community at the University of York. This study was approved by the University of York Psychology Department Ethics Committee.

### Stimuli

Stimuli were derived from images of a single male and a single female model from the NIMSTIM set (Tottenham et al., 2009). For these two models, closed mouth happy and neutral expression images were selected, and image-morphing techniques were used to create a sequence of images representing a morph-continuum between the two expressions. Following (Mayes et al., 2009) 45 fiducial points were used to identify corresponding spatial locations across the two images. These were placed at key locations on the face including the inner and outer canthi of the eyes, the centres of the pupils, multiple locations along the top and bottom of the upper and lower lips, and the face outline. Abrosoft Fantamorph (V 3.0) was then used to generate six intermediate images for each model, leading to a continuum of eight images for each model (2 original expressions, and 6 interpolated morphs between these). An oval frame was placed around each image to remove hair and background.

### Procedure

Participants viewed a series of trials that consisted of the presentation of a sequence of five images, in which each image was displayed for 517ms and then immediately replaced by the next image (0ms ISI). Stimuli were presented centrally, subtending a visual angle of approximately 3°. Sequences consisted of stepwise images from one of the morph-continua, either commencing with a relatively neutral image (Image 1 or Image 2 from the continuum) that was followed by three progressively more happy images, or commencing with a relatively happy image (Image 7 or Image 8) that was followed by three progressively less happy images. The fifth image in each sequence was then used to create Predictable or Unpredictable experimental conditions. In Predictable sequences the 5^th^ and final image conformed to the trajectory established by the preceding four images (either towards the full happy expression, or towards the neutral expression), whereas in Unpredictable sequences the 5^th^ and final image reversed the direction of the trajectory established by the four preceding images (for example, if the first four images were in increasing morphed steps toward happiness, the final image in an Unpredictable sequence would involve a morphed step back toward neutral).

By using both happy to neutral and neutral to happy sequences in the first 4 images, we were able to match the set of final image transitions across the Predictable and Unpredictable conditions. That is to say each possible penultimate-to–final image transition for a trial in the Predictable condition was matched to an identical penultimate-to–final image transition for an Unpredictable trial whose initial trajectory was in the opposite direction (see Figure 1). Thus, the set of Predictable trials consisted of image sequences 1-2-3-4-5, 2-3-4-5-6, 8-7-6-5-4 and 7-6-5-4-3, whilst the set of Unpredictable trials of image sequences was 1-2-3-4-3, 2-3-4-5-4, 8-7-6-5-6 and 7-6-5-4-5. This balancing of the fourth and fifth images in each sequence across conditions means that any differences in the ERPs to Predictable versus Unpredictable trials must arise as a consequence of the sequence of images preceding the final (fourth to fifth) image transition (i.e. the context), and cannot be due to any property of the final image transitions themselves (as exactly the same transitions were used in each condition).

**Figure 1.**
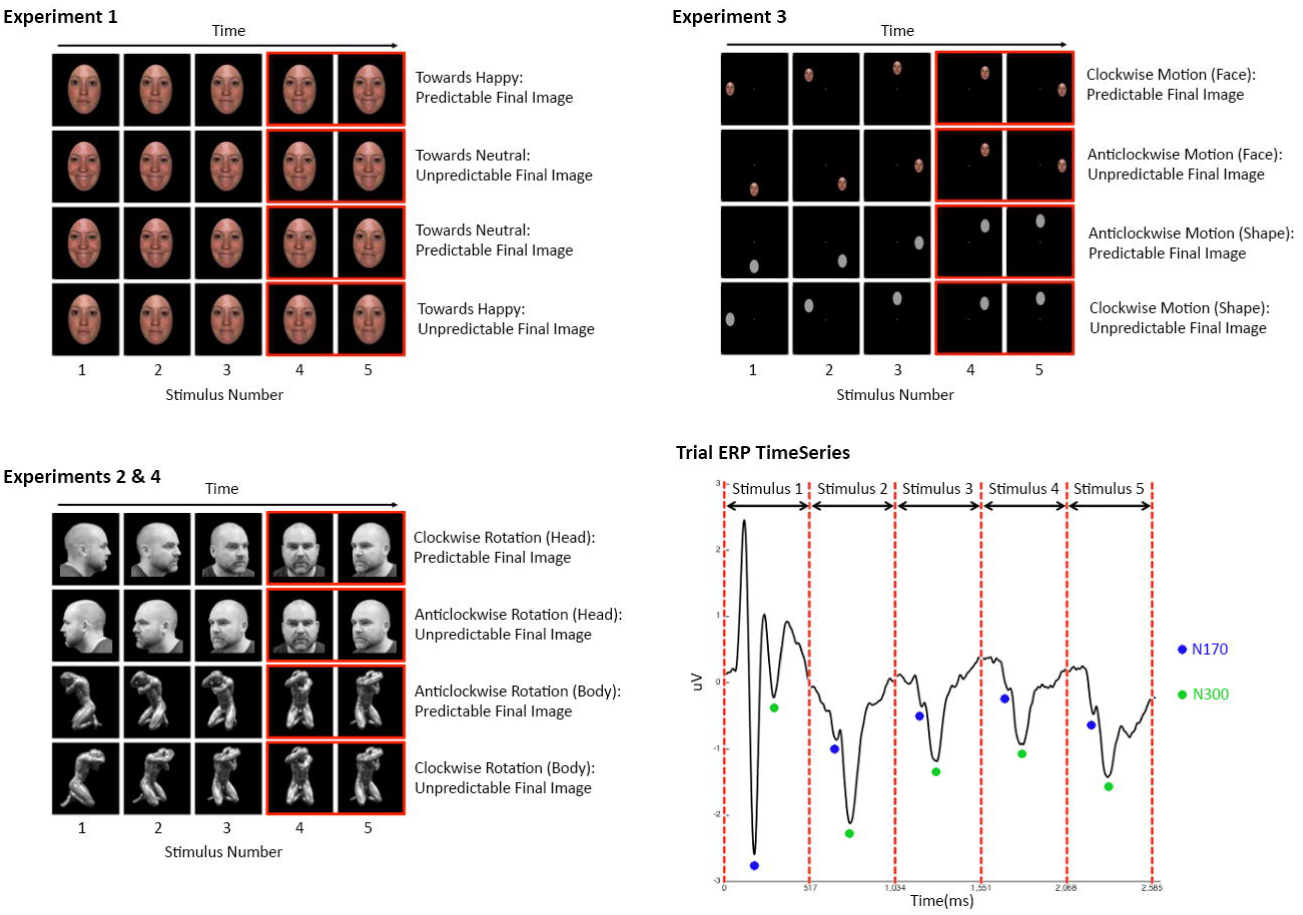
Examples of stimulus sequences used in Experiments 1-4. In each experiment the first four images in the sequence were used to create a regular change trajectory. In Experiment 1 (Top Left) this involved sequential steps along a morph continuum either from a more neutral face to a more happy face (Towards Happy), or from a more happy face to a more neutral face (Towards Neutral). In Experiments 2 and 4 (Bottom Left) sequences involved the clockwise or counterclockwise rigid-rotation of Heads or Bodies. In Experiment 3 (Top Right), stimulus sequences involved the locational rotation (clockwise or counterclockwise) around points corresponding to the main winds of the compass of either a Face or a Shape (grey oval). In all cases the initial stimulus was preceded by a fixation screen. Each of the five stimuli was presented for 517ms and was replaced by the next stimulus in the sequence with no intervening fixation screen (ISI 0ms). The final image in the sequence could be Predictable with respect to the trajectory established by the preceding images (i.e. the next image in the implied sequence) or Unpredictable (a step backward in the sequence). We were able to match across the set of penultimate-to-final image transitions (between the fourth and fifth images in each sequence, marked here with red frames) by pairing each Predictable trial sequence with an Unpredictable trial sequence whose initial trajectory was opposite to that of the Predictable sequence, but involved a reversal in the direction of the trajectory at the final step. This ensured that any differences observed between Unpredictable and Predictable final stimuli must be a consequence of the preceding events rather than the stimulus transition itself. A schematic waveform identifying ERP components in response to the sequential stimulus onsets across a trial is shown (Bottom Right). For all experiments analyses focus upon the ERP to the Predictable or Unpredictable 5^th^ stimulus.

It is important to note, that with a frame rate of <2 frames per second (fps) our stimuli are not perceived as fluid motion (which requires a minimum of 12fps) but as a series of still images. The transitions between images are “jumps”. The extent to which these “jumps” may be perceived as “motion” is post-hoc since the spatial translation of the stimuli implies that motion must have occurred.

There were equal numbers of Predictable and Unpredictable sequence trials (160 each), presented in a randomized order. Trials were separated by a 1017ms inter-trial interval during which a central fixation-cross was presented in an otherwise blank screen.

Participants were asked to maintain their gaze on the central fixation point. In order to maintain visual attention, they completed a simple vigilance task. This involved a set of 32 trials that were randomly interleaved with the main experimental trials. They were identical to the experimental trials with the exception that one of the images in the sequence included a small red-dot appearing at a location on the face close to the centre of the screen. This dot could appear (with approximately equal likelihood) on any of the five images constituting the trial. Participants were required to respond via a button press whenever they saw an image containing a red-dot. “Red-dot” trials were coded separately, and were not included in the analysis of ERP data. Apart from this red-dot monitoring task, the experiment involved passive viewing of the stimuli - participants were never asked anything about whether the sequences were predictable or unpredictable.

The task was delivered using Psychopy software (version 1.75) running on an Intel Pentium 4 HT computer, and the visual stimuli were presented on a 23” TFT LCD widescreen monitor with a 1340 × 1084 pixel resolution. Participants were seated approximately 60cm away from the screen.

### EEG Recording and Analyses

EEG was collected with a sampling rate of 1000Hz on 64 channels using an ANT ASAlab high-speed amplifier, from scalp sites corresponding to the extended International 10-20 electrode montage using a WaveGuard cap. An averaged reference was used and impedance values were kept below 20KOhms. Vertical and horizontal EOG measures were taken using bipolar electrode pairs placed above and below the left eye, and proximal to the outer canthus of each eye, as a basis for detecting and attenuating eye-movement artifacts in the EEG data post recording. EEG data were filtered using a bandpass filter (0.3-30Hz, slope 24dB per octave) with a notch-filter at 50Hz. Eye-movement artifacts associated with blinks were attenuated using the Gratton-Coles procedure (Gratton et al., 1983).

Data were segmented into epochs beginning 200ms before the onset of each trial and continuing for 2600ms to encompass the complete sequence of 5 images. Averaged ERPs were generated for each condition, time-locked to the onset of the first image in each trial. ERPs were baselined to the period between 150-0ms prior to the onset of the 5^th^ image in each sequence. For each participant and condition, N170 amplitudes were calculated as the average amplitude between 140ms-200ms following stimulus onset (for the 4^th^ and 5^th^ stimulus in the sequence), for electrodes P7 and P8.

### Experiment 2

We examined rigid rotational trajectories of images of heads and bodies. As before, trials consisted of the presentation of sequences of images that followed a consistent trajectory across the first four images, and where the final image either did or did not conform to the established pattern. We predicted that if the N170 is an index of predictive mechanisms, N170 amplitudes to Unpredictable final images would be larger than to Predictable final images, regardless of stimulus type.

### Participants

There were 20 participants (14 female; mean age 23.6 years, SD4.0). These participants were a different sample to those who had participated in the previous experiment. All were an opportunity sample from the undergraduate and postgraduate community at the University of York. This study was approved by the University of York Psychology Department Ethics Committee.

### Stimuli

All stimuli were acquired with a digital camera, captured at 7 different angles per stimulus, with a 30° angle of rotation separating each image. For the head images this included images starting with a side-on profile, facing to the left, rotating through steps of 30° through a frontal face image to a side-on profile facing right. Each category of stimulus used one male and one female exemplar, totaling four exemplars overall. The Head category subjects were Caucasian and their images were cropped at the neck. Body stimuli were created from statuettes representing human figures; where the face was visible it was blurred using a pixellation filter using the GIMP^TM^ software.

### Procedure

As in Experiment 1, each trial consisted of a sequence of 5 images, where the final image could be Predictable or Unpredictable with respect to the rotational trajectory established by the preceding images. Stimuli were presented centrally, subtending a visual angle of approximately 3°. Sequences depicted the clockwise (as viewed from above) or anticlockwise rotation (in 30° increments) of either heads or bodies. On Unpredictable trials the direction of rotation established across the first four images was reversed for the final image. As in Experiment 1, we were able to match the set of final image transitions across the Predictable and Unpredictable sequences (see Figure 1). There were 80 trials each with Predictable Heads, Unpredictable Heads, Predictable Bodies and Unpredictable Bodies. All trial types were randomly interleaved. Within trials, each image was presented for 517ms (0ms ISI), and there was an inter-trial interval of 1017ms. As for the previous experiment participants performed a “red-dot” vigilance task. There were a total of 64 “red-dot” trials (equal numbers of Head and Body trials). Again, participants were asked to maintain their gaze on the central fixation point. EEG recording, processing and analyses parameters were identical to those for Experiment 1.

### Experiment 3

This experiment used a simple 2D shape and a neutral face as stimuli. Since 2D shapes are not amenable to changes involving expression or 3D rotation, we used trajectories involving a series of step-wise changes in location around the initial fixation-point, with the final image in the sequence either in the expected location from the established sequence or a different location that involved a step back from the established direction. Again, we predicted that if the N170 is an index of predictive mechanisms, we should find increased N170 amplitudes to Unpredictable compared to Predictable final images, for both types of stimuli (face and shape).

### Participants

There were 18 participants (8 female; mean age 21.2 years, SD1.2). These participants were a different sample to those who had participated in the previous two experiments. All were an opportunity sample from the undergraduate and postgraduate community at the University of York. This study was approved by the University of York Psychology Department Ethics Committee.

### Stimuli

The stimuli were an image of a face, and an image of a grey oval. The face was a female (taken from Experiment 1) with a neutral expression, cropped within an oval frame and displayed on a black background. The grey oval was the same size as the oval frame displaying the face, and was also displayed on a black background.

### Procedure

Trials again consisted of sequences of 5 images. Stimuli were presented subtending approximately 1.5° of visual angle and offset by approximately 1.5° from the central fixation point (where a small grey dot was presented). The first stimulus in each trial appeared at one of the 8 main compass winds with respect to the central fixation point. The location of the first image was randomly selected, with each of the 8 locations occurring with equal frequency. Subsequent images were presented such that stimuli moved by one location around the compass winds in a consistent direction (clockwise or anticlockwise with equal frequency) across the first four images in each trial. For Predictable trials the final (fifth) image appeared in the location consistent with the trajectory established by the preceding four images. For Unpredictable trials the final image appeared in a location that reversed the established trajectory.

Within trials each stimulus was presented for 517ms and was replaced immediately by its successor (0ms ISI). There was an inter-trial interval of 1017ms. For each of the four conditions (Predictable Face, Unpredictable Face, Predictable Shape, and Unpredictable Shape) there were 80 trials, which occurred in a randomized order. As with the previous experiments we matched the set of final image transitions across the Predictable and Unpredictable trials. Examples of stimulus sequences are shown in Figure 6. Similarly to the previous experiments participants performed a “red-dot” vigilance task. There were a total of 64 “red-dot” trials (equal numbers of Face and Shape trials). Again, participants were asked to maintain their gaze on the central fixation point. EEG recording, processing and analyses parameters were identical to those for Experiment 1.

### Experiment 4

Here Experiment 2 was replicated using magnetoencephalography (MEG) to localize neural sources whose activity discriminated Unpredictable versus Predictable stimulus onsets in a time window consistent with the M170. In order to achieve this we used a recently developed beamformer metric - the Difference Stability Index (DSI) (Simpson et al., 2015). This metric is designed to identify locations in brain space whose evoked response most consistently differentiates two experimental conditions.

Since our paradigm involves trajectories of implied motion, it seems reasonable to conjecture that brain areas that are involved in motion perception might also be involved in generating predictions about the next stimulus in the context of the types of sequence that we present. We therefore hypothesized that an expectation violation signals consistent with the M170 latency would be localized to areas of the visual cortex involved in the processing of implied stimulus motion.

### Participants

There were a new sample of 20 participants (9 female; mean age 23.9 years SD5.4). All were an opportunity sample from the undergraduate and postgraduate community at the University of York. Two participants were excluded from the analysis, one due to a systematic blink artefact across trials, and one because of a large signal artefact during data acquisition, which made gaining a meaningful signal untenable. This study was approved by the York Neuroimaging Centre Ethics Committee.

### Stimuli and Procedure

Stimulus images and trial sequences were identical to those for Experiment 2. Images were projected using a Dukane 8942 ImagePro 4500 lumens LCD projector, projected onto a custom suspended 1.5 × 1.2 m fabric rear projection screen filling more than 65 × 30 degrees visual field in the MEG scanner. The stimuli subtended approximately 3° of visual angle.

### MEG Data Acquisition and Analysis

MEG Data were acquired using a 4D Neuroimaging Magnes 3600 system with 248 magnetometer sensors. The data were recorded at a rate of 678.17Hz, with an online 200Hz low pass filter for ~21 minutes. Three malfunctioning sensors were removed from the data analysis of all participants. Five reference location coils were used to monitor head position at the beginning and end of each recording. Movement of the five reference coils was limited to a threshold of 0.81cm. Each channel of the 320 epochs of data acquired for each participant was visually inspected for magnetic field fluctuations or physiological artifacts such as blinks, swallows, or movement. An average of 9.06 epochs (SD7.23) was rejected per individual. 1 Hz high pass and 40Hz low pass filters were applied to the data to improve the signal to noise ratio.

The location of 5 fiducial landmarks and a digital head shape were recorded prior to acquisition using a Polhemus Fastrack 3D digitiser. To enable anatomical inference in source-space, each individual’s digitised head shape was coregistered with an anatomical MRI scan using surface matching (Kozinska et al., 2001). A high-resolution T1-weighted structural MRI was acquired with a GE 3.0 T HDx Excite MRI scanner, using an 8 channel head coil and a sagittal isotropic 3D Fast Spoiled Gradient-Recalled Echo sequence. The spatial resolution of the scan was 1.13 × 1.13 × 1.0 mm, reconstructed to 1 mm isotropic resolution, with TR/TE/flip angle of 7.8 ms/3 ms/20 degrees. The field of view was 290 × 290 × 176, and in-plane resolution 256 × 256 × 176.

The source space analysis carried out for this work was based on a vectorised, linearly constrained minimum variance beamformer (Van Veen et al., 1997). The “weights” of the beamformer solution were calculated using equation *n*:

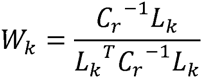

Where ***W*_*k*_** is the three dimensional weight vector for point *k*, ***L*_*k*_** is the three dimensional leadfield for point *k* and ***C*_*r*_** is the regularised estimate of the covariance. Here regularisation was applied using the smallest eigenvalue of ***C***.

The spatial beamformer relies upon analysis of the covariance structure across a set of trials. Because of this, temporal segments of the trials for which there is no discernible evoked signal hamper the determination of the set of weights that maximise the beamformer’s precision in inverting the sensor-level signal into source space. For this reason we defined a time-window that attempted to maximise the inclusion of time-points where brain signals are present whilst excluding time-points where brain signals are absent. To define this window of interest we calculated the Root Mean Square (RMS) signal amplitude across all sensors for each individual participant in the face condition, and then averaged these across all participants. From these data, we identified the RMS minima, which indicated the likely boundaries between evoked events. To define our analysis window, the RMS minimum associated with the first post-stimulus reversal was used as the start of our analysis window, and the first reversal below pre-stimulus RMS levels was used as the end of the window. We thus defined a time-window of 72ms-413ms post-stimulus onset for calculating covariance estimates. This window could then itself be separated into shorter time periods for more detailed analysis (see below), the overall window of 72ms-413ms is used here only to create the covariance estimates for the beamformer.

The beamformer weights, when applied to the recorded data, yield a three dimensional time series, or “virtual electrode” at each point in a source-space grid based on the MNI template, with grid-points separated by 5mm. These projected virtual electrodes allowed us to perform group-level analyses addressing the source localization of generators of the “expectancy violation N170” as indexed from our Unpredictable vs. Predictable trials. This was achieved using the beamformer metric known as the Difference Stability Index (DSI) first described by Simpson and colleagues (Simpson et al., 2015). In calculating the DSI metric, 3D virtual electrode (VE) time series are estimated for each source-space grid location for two different experimental conditions and a subtraction waveform is generated across the two conditions. The DSI searches through a set of 163 potential orientations to locate the orientation that maximizes the estimated stability of the phase-locked time course of the difference waveform. The estimate of signal stability is derived through a permutation method which estimates the average correlation across random split-halves of the set of trials. In essence it finds source space locations where there is greatest trial-by-trial consistency in the evoked brain response to a particular class of stimuli.

For group-level statistical inference we first generated a set of surrogate datasets in which any evoked activity was destroyed (through randomly sign-flipping 50% of the trials), and then performed a non-parametric label-permutation test to generate a null distribution of maximum pseudo-t statistics across the whole source-space grid. Group-level DSI values that exceeded the 99^th^ percentile of the null distribution were considered to be significant. It should be noted that by constructing null distributions based upon the maximum pseudo-t based upon permutation statistics across all of the grid-points within the brain volume, this method implicitly accounts for multiple comparisons across the entire set of tests (Nichols and Holmes, 2002).

We performed the DSI analysis localizing generators of the most stable differences in evoked responses to Unpredictable versus Predictable trails (across both Heads and Bodies) in response to the final image onset of the sequence. To localize the M170, this analysis considered signal stability across the period 110ms-210ms post-stimulus onset. This time-window centers around the expected M170 latency and is of sufficient duration to ensure stable estimates of measures of correlation coefficients. Whilst this choice of time-window does not exclude the possibility that the evoked brain signal at a particular location might differentiate between experimental conditions at other latencies outside this range, the choice of window is hypothesis-driven by our ERP findings and it maximises the likelihood that any stable differences in evoked responses to different conditions that are identified are attributable to differences in the M170 latency period.

## Results

For Experiment 1, one participant showed highly erratic ERPs that deviated substantially (i.e. by more than 3 standard deviations) from the other participants across a large portion of the trial (see Supplementary Materials). This participant was therefore excluded from analyses. No other participants were excluded from any of the other EEG experiments.

Grand Averaged ERPs across all participants for each of the conditions in Experiments 1-3 are shown in Figure 2. As can be seen, across these experiments a large P1-N170 complex was consistently evident to the onset of the first stimulus in a sequence as well as a later negative inflection of the ERP (the N300). Although P1 amplitudes were noticeably attenuated to subsequent stimuli in the sequence, there was a clearly observable (but attenuated) N170 to each subsequent stimulus, as well as an N300.

Importantly, and in line with our predictions, across all three experiments and conditions there were clearly observable modulations of the ERP to the fifth stimulus in the sequence as a function of stimulus predictability. Statistical treatment of these comparisons is detailed below.

### Experiment 1

We hypothesized that N170 amplitudes would be modulated by stimulus predictability. The critical comparison for testing our hypothesis involves N170 amplitudes in response to the final image in the sequence, comparing Predictable versus Unpredictable final images. However it is important to contextualize this comparison by showing that there were no differences between these conditions immediately prior to the final stimulus onset. We therefore performed a three-way repeated-measures ANOVA comparing N170 amplitudes to Predictable versus Unpredictable sequences in response to the onsets of the 4^th^ image in each sequence and in response to the 5^th^ image in each sequence, for electrodes P7 and P8 respectively. Planned contrasts were used to specifically test whether there were differences between Predictable and Unpredictable sequences in response to the 4^th^ stimulus (where there should be no differences between conditions) and in response to the 5^th^ stimulus (where we predict that differences between conditions should be apparent). The ANOVA showed that there was a main effect of sequential step (4^th^ versus 5^th^) (F(1,17)=5.97, p=.026, ns, *partial eta squared*=.260), and a significant main effect of stimulus predictability (F(1,17)=7.52, p=.014, *partial eta squared*=.307), and a main effect of laterality (electrode P7 versus P8) (F(1,17)=9.67, p=.006, *partial eta squared*=.363). Importantly there was a significant sequence step by stimulus predictability interaction (F(1,17)=26.08, p<.001, *partial eta squared*=.605), but there were no other significant interactions.

In line with our hypotheses, planned contrasts revealed that there were no differences in N170 amplitudes between Predictable and Unpredictable sequences in response to the 4^th^ stimulus onset (F(1,17)=0.06, p=.806, *ns*, *partial eta squared*=.004) but there were significant differences between conditions in response to the 5^th^ stimulus onset (F(1,17)=19.53, p<.001, *partial eta squared*=.535).

Visual inspection of the Grand Averaged ERP waveforms to Predictable versus Unpredictable final images (see Figure 2) suggested later latency differences to also exist between these conditions. As a supplementary analysis we extracted ERP amplitudes for the N300 component between 200-300ms following the 5^th^ stimulus onset. A repeated measures ANOVA was performed comparing Predictable to Unpredictable trials, showing that the N300 amplitude was larger to Unpredictable trials (F(1,17)=22.53, p<.001, *partial eta squared*=.570). However, when the N300 amplitude was recalculated as a difference from the N170 amplitude (by subtracting the N170 amplitude from the N300 amplitude), there were no significant differences between conditions (F(1,17)=0.17, p=.687, *ns*, *partial eta squared*=.010), implying that the differences observed at the N300 were likely to be a follow-on consequence of the differences at N170.

**Figure 2.**
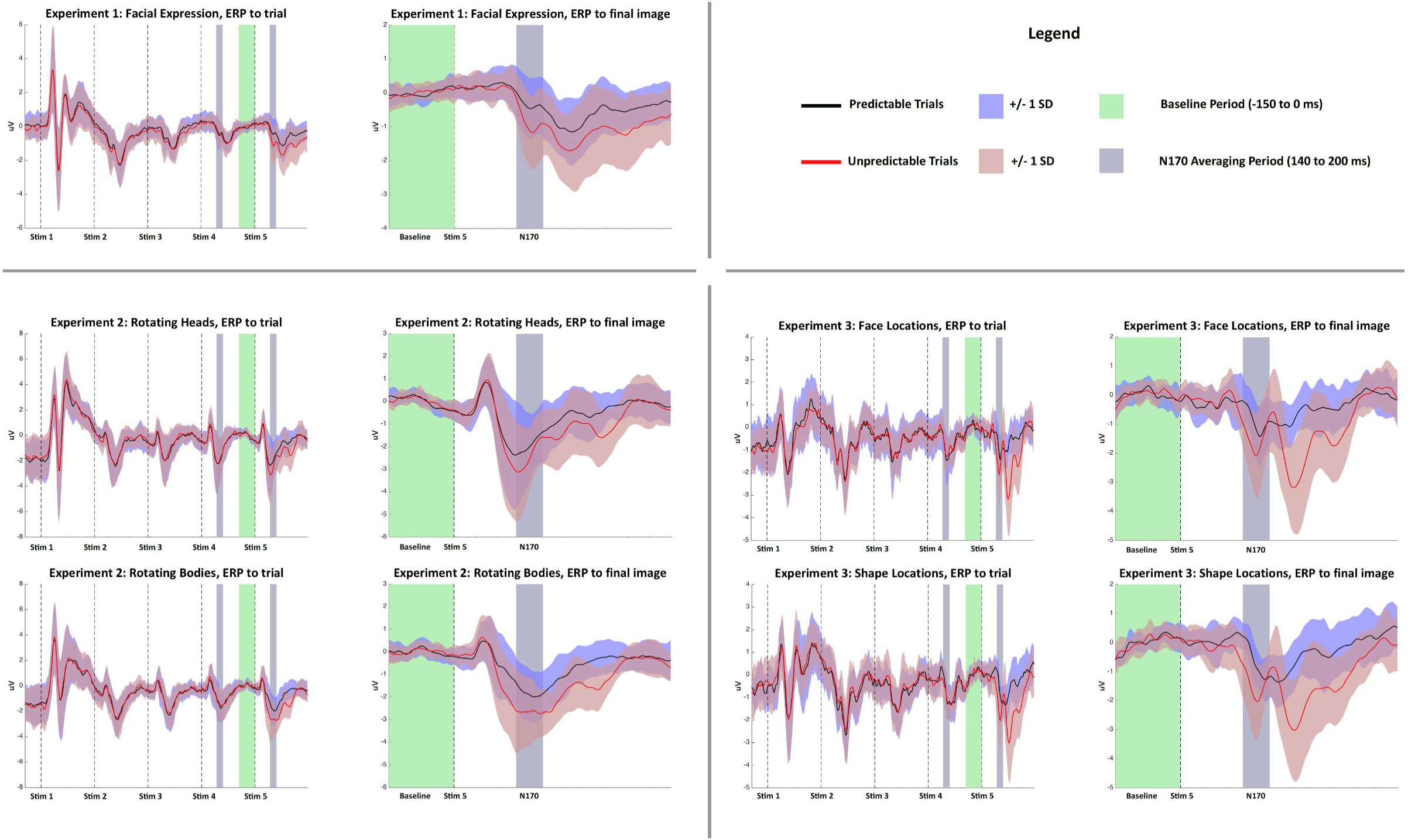
Grand Averaged ERPs across all participants to Unpredictable sequences (red) and Predictable sequences (black) for Experiments 1-3. Left column images show ERPs across the whole trial sequence of five consecutive images, whilst Right column images focus upon ERPs to the critical 5^th^ image onset.

### Experiment 2

We hypothesized that N170 amplitudes would be modulated by stimulus predictability, and that this should happen regardless of stimulus type. As in Experiment 1, the critical comparison is between N170 amplitudes in response to Predictable and Unpredictable final images. Again it was important to show that there were no differences between these conditions immediately prior to the final stimulus onset. We therefore performed a four-way repeated-measures ANOVA with factors of laterality (electrode P7 vs. P8), stimulus type (Heads vs. Bodies) sequential step (4^th^ vs. 5 ^th^ image onset) and sequence predictability (Predictable vs. Unpredictable). Planned contrasts were employed to test specific hypotheses that there should be no differences between N170 amplitudes to Predictable versus Unpredictable sequences in response to the 4^th^ stimulus onset for either Head or Body stimuli, but that such differences should occur in response to the 5^th^ stimulus onset for both types of stimulus object.

The ANOVA revealed no main effect of stimulus type (F(1,19)=1.30, p=.268, ns, *partial eta squared*=.064). There were however significant main effects of laterality (F(1,19)=11.55, p=.003, *partial eta squared*=.378), sequential step (F(1,19)=17.43, p=.001, *partial eta squared*=.478), and stimulus predictability (F(1,19)=12.15, p=.002, *partial eta squared*=.390), as well as significant interactions of stimulus type by sequential step (F(1,19)=4.72, p=.037, *partial eta squared*=.210), and laterality by predictability (F(1,19)=5.20, p=.034, *partial eta squared*=.215)

Most importantly, there was an interaction of sequential step by stimulus predictability (F(1,19)=17.92, p=.001, *partial eta squared*=.485). This interaction was not modulated by effects of stimulus type or laterality (i.e. there were no three-way or four-way interactions). In line with our hypotheses, planned comparisons revealed that for both Heads (F(1,19)=6.07, p=.023, *partial eta squared*=.242) and Bodies (F(1,19)=27.56, p<.001, *partial eta squared*=.592) there were greater N170 amplitudes to Unpredictable than to Predictable sequences in response to the 5^th^ stimulus onset. Crucially, there were no differences in N170 amplitudes to 4^th^ image onsets between Predictable and Unpredictable sequences for either Heads (F(1,19)=0.30, p=.589, *ns, partial eta squared*=.016) or Bodies (F(1,19)=0.02, p=.897, *ns, partial eta squared*=.001).

Given the extensive literature demonstrating greater N170 amplitudes to faces than to other stimulus types, it might be considered slightly surprising that there was no main effect of stimulus type across the 4^th^ and 5^th^ images in the stimulus sequences. Since our research paradigm is novel, and without clear analogues in the existing literature, there are a range of reasons why this might have been the case that will require further investigation. Although this question is not directly relevant to the current research, in order to link our data to the existing literature we performed a supplementary analysis examining N170 amplitudes to the first occurring stimulus in each type of stimulus sequence. This analysis revealed that initial N170 amplitudes to the first stimulus onset were greater to Heads (mean: −1.069 uV, SE: 0.650 uV) than to Bodies (mean: 0.013 uV, SE: 0.575 uV) (F(1,19)=17.93, p<.001, *partial eta squared*=.486).

As with Experiment 1, inspection of the Grand Averaged ERP waveforms (see Figure 2) suggested later latency differences also to exist between Predictable and Unpredictable trials. As a supplementary analysis we extracted ERP amplitudes for the N300 component between 200-300ms following the 5^th^ stimulus onset. Looking at raw ERP scores there was a difference in N300 between Predictable and Unpredictable final images (F(1,19)=6.18, p=.022, *partial eta squared*=.245). However, when the N300 amplitude was recalculated as a difference from the N170 amplitude (by subtracting the N170 amplitude from the N300 amplitude), there were no significant differences between conditions (F(1,19)=0.63, p=.437, *ns*, *partial eta squared*=.032). As with Experiment 1, this implies that the differences observed at the N300 were likely to be a follow-on consequence of the earlier differences at N170.

### Experiment 3

Following the same strategy as Experiment 2 we performed four-way repeated-measures ANOVA with factors of laterality (electrode P7 vs. P8), stimulus type (Faces vs. Shapes) sequential step (4^th^ vs. 5^th^ image onset) and sequence predictability (Predictable vs. Unpredictable), with planned contrasts comparing N170 amplitudes to Predictable and Unpredictable sequences in response to the 4^th^ stimulus onset, and 5^th^ stimulus onset, for Faces and for Shapes. The ANOVA showed no main effect of laterality F(1,17)=0.01, p=.930, *ns*, *partial eta squared*=.001), or stimulus type F(1,17)=1.65 p=.216, *ns*, *partial eta squared*=.089), or sequential step (F(1,17)=3.33, p=.066, *partial eta squared*=.185). There was a significant main effect of stimulus predictability (F(1,17)=7.731, p=.013, *partial eta squared*=.313), as well as a significant interaction of sequential step by stimulus predictability (F(1,17)=10.58, p=.005, *partial eta squared*=.384). There were no other significant interactions.

In line with our hypotheses, planned comparisons revealed that for both Faces (F(1,17)=6.52, p=.021, *partial eta squared*=.277) and Shapes (F(1,17)=13.64, p=.002, *partial eta squared*=.445) there were greater N170 amplitudes to Unpredictable than to Predictable sequences in response to the 5^th^ stimulus onset. Importantly, there were no differences in N170 amplitudes to 4^th^ image onsets between Predictable and Unpredictable sequences for either Faces (F(1,17)=0.20, p=.663, *ns, partial eta squared*=.011) or Shapes (F(1,17)=0.28, p=.602, *ns*, *partial eta squared*=.016).

As with the previous experiments supplementary analyses were performed to examine later latency effects of stimulus predictability. We extracted ERP amplitudes for the N300 component between 200-300ms following the 5^th^ stimulus onset. Looking at raw ERP scores there was a significant difference between Predictable and Unpredictable trials (F(1,17)=37.68, p<.001, *partial eta squared*=.689). When these were recalculated as differences from the N170 amplitude (by subtracting the N170 amplitude from the N300 amplitude), there remained a significant difference between Predictable and Unpredictable final images (F(1,17)=6.40, p=.022, *partial eta squared*=.274).

### Experiment 4

DSI values were generated comparing the evoked responses to Unpredictable versus Predictable Head or Body stimuli across a time-window (110ms-210ms) consistent with the M170. These analyses revealed the strongest statistically significant differences in stable evoked responses to Unpredictable versus Predictable stimuli in areas of the occipito-temporal cortices known to be involved in the motion and implied motion of objects and biological agents including the right Middle Temporal gyrus (MT) and Superior Temporal Sulcus (STS) (see Figure 3). There were further statistically significant sources where the evoked response consistently differentiated Unpredictable versus Unpredictable stimuli in the right Angular gyrus, the right Superior Parietal Lobule, right Central and Parietal Opercular cortices, and the left posterior Cingulate Gyrus. MNI coordinates of peak DSI values are reported in Table 3. A Virtual Electrode showing estimated Grand Averaged Event Related Field magnitudes across all participants to Predictable and Unpredictable trials is shown (Figure 3) for MNI coordinates corresponding to the peak DSI value in the right Middle Temporal Gyrus (MT: 53,-56,13). This VE clearly demonstrates greater magnitude responses to Unpredictable versus Predictable trials following the final stimulus onset, that occur with a latency consistent with N/M170.

**Figure 3.**
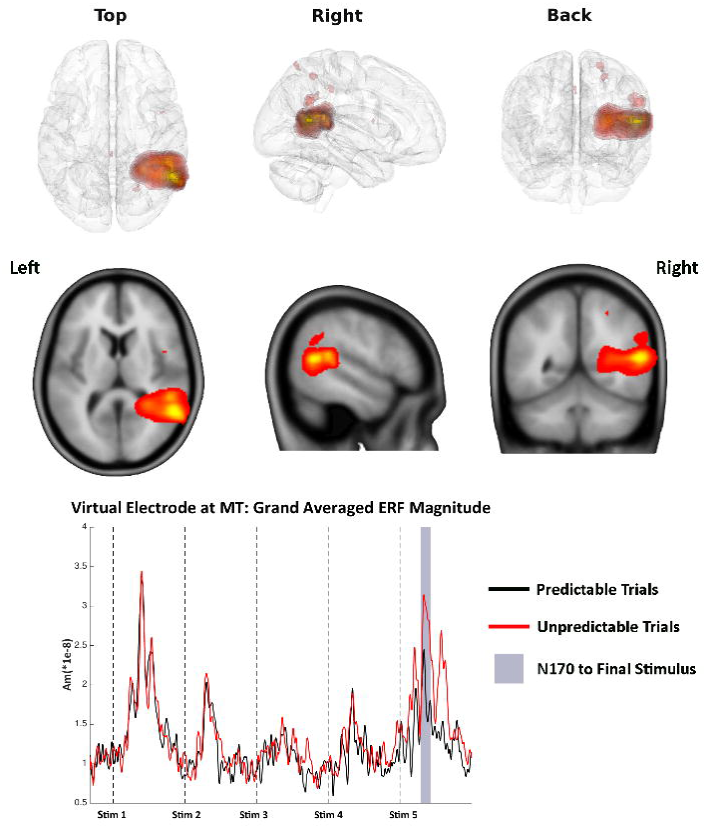
Experiment 4: MEG DSI source localisation of expectancy violation signals to Unpredictable versus Predictable stimuli across the M170 time-window. Images generated using the DataViewer3D software (Gouws et al., 2009).

**Table 1.** For Experiments 1-3, Mean ERP Amplitudes (Uv) (Standard Errors in brackets) for the N170 to the 4^th^ and 5^th^ stimulus onset and the N300 to the 5^th^ stimulus onset by trial type (pooled across electrodes P7/P8).

|  | **N170 to 4<sup>th</sup>**<br>**Stimulus**<br>**P7/P8** | **N170 to 5<sup>th</sup>**<br>**Stimulus** | **N300 to 5<sup>th</sup>**<br>**Stimulus** |
| --- | --- | --- | --- |
| **Experiment 1** |  |  |  |
| Predictable | -0.402 (0.161) | -0.332 (0.170) | -0.918 (0.193) |
| Unpredictable | -0.379 (0.181) | -0.870 (0.222) | -1.408 (0.241) |
| **Experiment 2** |  |  |  |
| Predictable Head | -1.905 (0.375) | -1.927 (0.320) | -0.753 (0.229) |
| Unpredictable Head | -1.835 (0.375) | -2.436 (0.320) | -1.044 (0.184) |
| Predictable Body | -1.451 (0.277) | -1.822 (0.245) | -1.285 (0.276) |
| Unpredictable Body | -1.433 (0.270) | -2.658 (0.292) | -1.917 (0.195) |
| **Experiment 3** |  |  |  |
| Predictable Face | -1.027 (0.203) | -0.872 (0.169) | -0.776 (0.122) |
| Unpredictable Face | -0.946 (0.149) | -1.493 (0.240) | -2.041 (0.256) |
| Predictable Shape | -0.778 (0.144) | -0.657 (0.128) | -0.822 (0.152) |
| Unpredictable Shape | -0.872 (0.166) | -1.461 (0.235) | -2.053 (0.316) |

**Table 2.** Locations of peak DSI values for the contrast Unpredictable versus Predictable trials.

| Brain Area | MNI Coordinates | DSI-value | P value |
| --- | --- | --- | --- |
| R. Middle Temporal Gyrus | 53,-56,13 | 23.34 | < .001 |
| R. Superior Temporal Sulcus | 48,-41,18 | 18.91 | < .001 |
| R. Angular Gyrus | 53,-56,28 | 10.44 | < .01 |
| R. Superior Parietal Lobule | 23,-66,63 | 8.88 | < .01 |
| R. Cental Opercular Cortex | 43,4,13 | 8.22 | < .01 |
| L. Cingulate Gyrus | -2,-36,43 | 8.12 | < .01 |
| R. Parietal Operculum | 38,-21,18 | 7.79 | < .01 |

## Discussion

In three EEG experiments we demonstrated robust patterns of modulation of N170 amplitudes in which the N170 was greater to Unpredictable than to Predictable final image onsets (in the absence of such differences to the penultimate stimulus), across a range of stimulus types and contextual trajectories. In a further MEG experiment we localised the expectancy violation signal (with respect to rotational trajectories of Heads and Bodies) to brain areas MT and STS. In all of these experiments the matching of final image transitions across Predictable and Unpredictable trials means that these differences in N170 must be attributable to the sequence of preceding events.

In Experiment 1, N170 amplitudes were larger to Unpredictable expressions than to Predictable ones. We therefore suggest that enhanced N170 amplitudes to Unpredictable stimuli reflect violation of expectations concerning the “expression trajectory” established by the preceding images. The conclusion that some form of perceptual prediction is involved is therefore strongly motivated.

In Experiment 2 Unpredictable steps in rigid rotational trajectories (of Heads and Bodies) elicited a larger N170 response than Predictable ones. There was no main effect of stimulus type, and no interaction of stimulus type with predictability. This result is consistent with our suggestion that N170 can index prediction/error-checking mechanisms, and extends the findings of Experiment 1 to a different type of contextual trajectory (rotation) and to bodies and heads as well as faces. These findings suggest general mechanisms indexing visual prediction errors with respect to different types of motion trajectory.

Although there were no differences in N170 amplitudes between Heads and Bodies in response to the 4^th^ and 5^th^ occurring stimuli in the trial sequences of Experiment 2, differences did occur in response to the first occurring stimuli following the intertrial interval, such that the N170 was larger to Heads than to Bodies. This is consistent with previous literature (Liu et al., 2000; Rossion and Jacques, 2008; Eimer, 2011) showing an enhanced N170 response to faces compared to other objects, believed to index face specific perceptual processes. It is unclear why there were no differences between stimulus categories at the later stages of the trial, however, it is possible that this may reflect differential stimulus adaptation patterns (Simpson et al., 2015). Notwithstanding, the current results clearly demonstrate a robust modulation of the N170 by expectancy violations irrespective of whether those expecatations related to Heads or Bodies.

Since privileged status has been claimed for bodies as well as faces (Hietanen and Nummenmaa, 2011; Alho et al., 2015), we performed a further experiment involving simple shapes as stimuli for which any privileged status is unambiguously not the case. Thus, Experiment 3 considered location trajectories to both Face and simple Shape stimuli. As before, there was an increased N170 to Unpredicted final stimuli, irrespective of stimulus type. Again, our careful experimental control means that the observed effect must be attributable to the predictability of the final stimulus based on the sequence of images preceding the final image onset.

Across all these experiments, an alternative explanation for the findings might be that in Unpredictable sequences the 5^th^ image in the sequence was identical to the 3^rd^ image, whereas in the Predictable sequences each image appeared only once. In the fMRI literature there is ample evidence of both signal enhancement and signal suppression occurring as a consequence of stimulus repetition in different tasks and circumstances (see Segaert et al 2013 for a review). Reductions in repetition suppression to unexpected stimuli suggest that repetition suppression may be related to prediction error-minimisation (Summerfield et al, 2008). In EEG, a number of studies report increased power in certain frequency bands as a consequence of stimulus repetition (ie. Gruber & Müller, 2002; Gruber & Müller, 2005), however, these same studies report decreased ERP amplitudes in response to stimulus repeats. Repetition enhancement effects on the ERP amplitude have been (Morel et al, 2009), however, unlike the current experiments, stimulus repeats occurred at a latency of greater than one minute. Most studies reporting effects of short-latency stimulus repeats on the N/M170 report *reductions* in N/M170 amplitude to stimulus repetition (Kloth et al., 2010; Mercure et al., 2011; Eimer et al., 2011; Fu et al., 2012; Engell and McCarthy, 2014; Caharel et al., 2015; Cao et al., 2015; Feuerriegel et al., 2015; Simpson et al., 2015; Tian et al., 2015). That said, on the basis of the current experiments we cannot rule out the possibility that image repetitions may have had some influence on the ERP, since there is some evidence in the (auditory) ERP literature that repetition suppression and expectation suppression may interact (Todorovic & de Lange, 2012). In order to rule out such effects we are following up the current studies with a variant of the paradigm whose design avoids within trial image repetitions.

In the final Experiment we used MEG to examine the source generators of expectancy violation signals (to rigid rotations of Heads and Bodies) consistent with the M170 time-window. This revealed that the major sources of the expectancy violation signal for this time-window were localized to MT and STS.

This localization is interesting because an extensive literature points to a role for areas MT and STS in the processing of the motion (and implied motion) of objects and biological agents (Dubner and Zeki, 1971; Maunsell and Van Essen, 1983; Newsome and Pare, 1988; Allison et al., 2000; Kourtzi and Kanwisher, 2000; Johnston et al., 2013). Since our experiments focus upon expectations (and expectation violations) with respect to implied trajectories across a range of stimuli (facial expressions, heads, bodies, shapes), we find this localization of expectancy violation signals to these brain areas to be compelling. We suggest that predictive representations are generated with reference to ongoing context and tested against incoming stimulus attributes in brain substrates involved in processing biological stimulus motion. The increased M/N170 to Unpredictable final stimuli in our experiments then results from a mismatch between expected and actual inputs to these systems.

Importantly MT and STS are regions that are spatially distinct from the generators of the face-sensitive N/M170, which are most commonly reported as the fusiform and lingual gyri (Halgren et al., 2000; Gao et al., 2013; Perry and Singh, 2014; Simpson et al., 2015). This localisation of the classic face-sensitive N/M170 also argues against the observed effects in our experiments being attributable to stimulus repetition, since our MT and STS generators are not consistent with generators where stimulus repetition effects on the M170 have previously been demonstrated. Instead, our previous work (Simpson et al., 2015) reported reduced M170 amplitudes in the fusiform gyrus to repeated presentation of faces, and at the occipital pole for repetitions of both faces and objects. There were no M170 amplitude increases due to repetition of either type of stimuli.

More generally, then, it seems that the N/M170 can be localized to different sources with different paradigms. We therefore propose that the N/M170 signal is generated across widespread areas of the visual brain (see Simpson et al., 2015), and indexes processes relating to the resolution of stimulus driven and top-down (predictive) perceptual influences across a range of stimulus types and attributes. Here, we have compellingly demonstrated its relation to one such process, predictive coding. Having shown this, though, we do not dispute that other paradigms can track other influences on the N/M170, and this is consistent with the different source localizations observed.

That the MEG source localization detected exclusively right lateralized generators of N/M170 expectancy violation signals was unexpected, given the lack of laterality effects in the EEG data, and is worthy of comment and further exploration. Although in EEG Experiment 1 and Experiment 2 we reported main effects of laterality (such that N170 responses were greater at right lateralized electrode P8 than at P7), these effects did not interact with stimulus predictability. Similarly in Experiment 3, there was no interaction of laterality with stimulus predictability. Thus, all the EEG experiments point towards bilateral generators of the expectancy violation signal. We believe that this inconsistency in findings in our MEG versus EEG studies may be due to the nature of the DSI beamformer metric. The method is not based upon the comparison of signals amplitudes across different conditions, but rather in detecting differences in the trial-by-trial stability, or consistency, of the evoked signal (Hymers et al, 2010; Simpson et al, 2015).

In the current study a high DSI value indicates that there is a highly consistent difference (at the trial-by-trial level) between Predictable and Unpredictable trials. An ERP on the other hand, is a measure of signal the average evoked signal *amplitude* across a set of trials. Sets of trials with rather different characteristics could end up with similar ERPs as a function of the averaging process – for instance a “smaller” but more (trial-by-trial) consistent signal might end with a similar ERP amplitude to a “larger” but less (trial-by-trial) consistent signal. However, these signals would lead to very different DSI values – the small consistent signal would lead to a strong DSI, whereas the larger less consistent would lead to a much smaller DSI value. In our current MEG data set, the DSI metric clearly indicates highly *consistent differences* between Predictable versus Unpredictable trials in the right MT and STS. However, it does not show such differences in the left hemisphere despite our EEG data supporting bilateral generators of the expectancy violation signal. One possible explanation for this is that whilst left lateralized cortical regions are generating expectancy violation signals (that are detectable in the Grand Averaged ERP) that there is a greater trial-by-trial variability in these signals that renders them less detectable by the DSI beamformer metric. This will be an important issue to explore in future studies.

Across Experiments 1-3 our results also showed a later effect of stimulus predictability between 200-300ms, although in both Experiment 1 and Experiment 2 this could be accounted for simply as a follow-on effect resulting from earlier differences at N170. Previous literature has reported an expectation violation N300 ERP to gaze-shifts and arrows (Senju et al., 2006; Tipples et al., 2013). That these earlier studies showed no effects at N170 may reflect differences in analyses (both studies pooled across electrodes including some more anterior and parietal than those reported here) or to effects in these studies involving shifting the locus of visual attention rather than “expectancy violations” *per se*. A minor discrepancy with the previous experiments was that in Experiment 3 effects observed at the later N300 could not be accounted for simply as a follow-on effect from the N170. It is not clear why this was so, but we speculate that it may relate to previously reported N300 effects (Senju et al., 2006; Tipples et al., 2013). Similarly to those studies (and unlike our preceding experiments) trials in Experiment 3 demanded a shift in the locus of visual attention for each stimulus. Nonetheless, the existence of these later effects does not decrease our confidence that the N170 partly indexes expectation violations.

Although the current work is conceptually related to research relating to another electrophysiological signal that has also been suggested to index predictive mechanisms – the visual Mismatch Negativity (vMMN) - (Stefanics et al., 2014) there are important differences that distinguish our work from the vMMN. The vMMN is a difference waveform derived through subtracting the ERP response to a rare “deviant” stimulus from that to a common “standard” stimulus. Historically, vMMN studies have relied upon simple visual stimuli (Pazo-Alvarez et al., 2003). Although more recent studies have considered more complex stimuli including emotional faces (Zhao and Li, 2006; Chang et al., 2010) these have generally adopted a frequent standard verus rare deviant stimulation schedule. Where effects of sequences have been considered (Kimora et al, 2010) this has been as a function of the large-scale periodicity of the deviant stimulus. This is markedly different from the current experimental paradigm in which Predictable and Unpredictable final stimuli are equally frequent and expectations are established on the basis of the narrative trajectory of a short stimulus sequence.

We have shown that the N/M170 is strongly modulated by violations of expectations across different contextual trajectories and stimulus types. Moreover, we found no evidence that this modulation of the N/M170 by expectation was influenced by stimulus category. We propose therefore that the N1/M70 can index quite general processes relating to perceptual prediction and error-checking/resolution in the visual domain. From a more general perspective, predictive mechanisms might exist in part to maintain perceptual representations across naturally occurring interruptions of visual input. Although our experience of the visual world appears seamless and cinematic, in fact there are frequent breaks in visual input. The average person blinks around 15 times per minute, and for each blink visual input is blocked for around 200-250ms (Johns et al., 2009). Similarly, visual input is suppressed whenever we make a saccadic eye movement (Johns et al., 2009). Yet although visual inputs are frequently disrupted we are not aware of these breaks; our visual system involves mechanisms that can edit these gaps from our awareness. Predictive coding offers a viable model for how this may be accomplished, since ongoing predictions could form bridging representations spanning brief periods. However, the constructive nature of these representations may be a source of perceptual errors. We believe that the N/M170 may provide an invaluable tool for interrogating such processes.

In conclusion, we have demonstrated that the N/M170 is strongly modulated by expectation violations. This new insight has profound implications for the N/M170's future potential as a tool for understanding how the brain encodes and represents context in order to generate perceptual predictions, and how this may contribute to error-proneness in a range of settings and clinical conditions.

## Acknowledgements

We thank Philip Dwerryhouse and Junior Whitely for helping with stimuli and data collection. We also gratefully acknowledge the gentle encouragement of The Oily Rag Foundation (EN10002).

## References

Alho J, Salminen N, Sams M, Hietanen JK, Nummenmaa L (2015) Facilitated early cortical processing of nude human bodies. Biological Psychology 109:103–110.

Allison T, Puce A, McCarthy G (2000) Social perception from visual cues: Role of the STS region. Trends in Cognitive Sciences 4:267–278.

Bentin S, Allison T, Puce A, Perez E, McCarthy G (1996) Electrophysiological studies of face perception in humans. Journal of Cognitive Neuroscience 8:551–565.

Caharel S, Collet K, Rossion B (2015) The early visual encoding of a face (N170) is viewpoint-dependent: A parametric ERP-adaptation study. Biological Psychology 106:18–27.

Cao X, Ma X, Qi C (2015) N170 adaptation effect for repeated faces and words. Neuroscience 294:21–28.

Chang Y, Xu J, Shi N, Zhang B, Zhao L (2010) Dysfunction of processing task-irrelevant emotional faces in major depressive disorder patients revealed by expression-related visual MMN. Neuroscience Letters 472:33–37.

Dubner R, Zeki SM (1971) Response properties and receptive fields of cells in an anatomically defined region of the superior temporal sulcus in the monkey. Brain Research 35:528–532.

Eimer M (2011) The face-sensitive N170 component of the event-related brain potential. In: The Oxford Handbook of Face Perception (Calder A, Rhodes G, Johnson M, Haxby JV, eds), pp 329–344: Oxford University Press.

Eimer M, Gosling A, Nicholas S, Kiss M (2011) The N170 component and its links to configural face processing: A rapid neural adaptation study. Brain Research 1376:76–87.

Engell AD, McCarthy G (2014) Repetition suppression of face–selective evoked and induced EEG recorded from human cortex. Human Brain Mapping 35:4155–4162.

Feuerriegel D, Churches OF, Keage HA (2015) Is neural adaptation of the N170 category-specific? Effects of adaptor stimulus duration and interstimulus interval. International Journal of Psychophysiology 96:8–15.

Friston K, Kiebel S (2009) Cortical circuits for perceptual inference. Neural Networks 22:1093–1104.

Fu S, Feng C, Guo S, Luo Y, Parasuraman R (2012) Neural adaptation provides evidence for categorical differences in processing of faces and Chinese characters: An ERP study of the N170. PLoS One 7:e41103.

Furl N, van Rijsbergen NJ, Treves A, Friston KJ, Dolan RJ (2007) Experience-dependent coding of facial expression in superior temporal sulcus. Proceedings of the National Academy of Sciences 104:13485–13489.

Gao Z, Goldstein A, Harpaz Y, Hansel M, Zion - Golumbic E, Bentin S (2013) A magnetoencephalographic study of face processing: M170, gamma - band oscillations and source localization. Human Brain Mapping 34:1783–1795.

Gouws A, Woods W, Millman R, Morland A, Green G (2009) DataViewer3D: An open-source, cross-platform multi-modal neuroimaging data visualization tool. Frontiers in Neuroinformatics 3:9.

Gratton G, Coles MG, Donchin E (1983) A new method for off-line removal of ocular artifact. Electroencephalography and Clinical Neurophysiology 55:468–484.

Gregory RL (1980) Perceptions as hypotheses. Philosophical Transactions of the Royal Society of London B, Biological Sciences 290:181–197.

Gruber T, Müller MM (2002). Effects of picture repetition on induced gamma band responses, evoked potentials, and phase synchrony in the human EEG. Cognitive Brain Research, 13(3), 377–392.

Gruber T, Müller MM (2005). Oscillatory brain activity dissociates between associative stimulus content in a repetition priming task in the human EEG. Cerebral Cortex, 15(1), 109–116.

Halgren E, Raij T, Marinkovic K, Jousmäki V, Hari R (2000) Cognitive response profile of the human fusiform face area as determined by MEG. Cerebral Cortex 10:69–81.

Hietanen JK, Nummenmaa L (2011) The naked truth: The face and body sensitive N170 response is enhanced for nude bodies. PLoS One 6:e24408.

Hietanen JK, Astikainen P (2013) N170 response to facial expressions is modulated by the affective congruency between the emotional expression and preceding affective picture. Biological Psychology 92:114–124.

Hubbard TL (2005) Representational momentum and related displacements in spatial memory: A review of the findings. Psychonomic Bulletin & Review 12:822–851.

Hymers M, Prendergast G, Johnson SR, Green GGR (2010) Source Stability Indext: A novel beamforming based localisation metric. Neuroimage 49, 1385–1397.

Johns M, Crowley K, Chapman R, Tucker A, Hocking C (2009) The effect of blinks and saccadic eye movements on visual reaction times. Attention, Perception, & Psychophysics 71:783–788.

Johnston P, Molyneux R, Young AW (2015) The N170 observed “in the wild”: Robust event-related potentials to faces in cluttered dynamic visual scenes. Social Cognitive and Affective Neuroscience 10.

Johnston P, Mayes A, Hughes M, Young AW (2013) Brain networks subserving the evaluation of static and dynamic facial expressions. Cortex 49:2462–2472.

Kaufman J, Johnston P (2014) Facial motion engages predictive visual mechanisms. PLoS One 9:e91037.

Kloth N, Schweinberger SR, Kovács G (2010) Neural correlates of generic versus gender-specific face adaptation. Journal of Cognitive Neuroscience 22:2345–2356.

Kourtzi Z, Kanwisher N (2000) Activation in human MT/MST by static images with implied motion. Journal of Cognitive Neuroscience 12:48–55.

Kozinska D, Carducci F, Nowinski K (2001) Automatic alignment of EEG/MEG and MRI data sets. Clinical Neurophysiology 112:1553–1561.

Levita L, Howsley P, Jordan J, Johnston P (2015) Potentiation of the early visual response to learned danger signals in adults and adolescents. Social Cognitive and Affective Neuroscience 10:269–277.

Liu J, Higuchi M, Marantz A, Kanwisher N (2000) The selectivity of the occipitotemporal M170 for faces. NeuroReport 11.

Maunsell JH, Van Essen DC (1983) Functional properties of neurons in middle temporal visual area of the macaque monkey. I. Selectivity for stimulus direction, speed, and orientation. Journal of Neurophysiology 49:1127–1147.

Mayes AK, Pipingas A, Silberstein RB, Johnston P (2009) Steady state visually evoked potential correlates of static and dynamic emotional face processing. Brain Topography 22:145–157.

McCandliss BD, Cohen L, Dehaene S (2003) The visual word form area: Expertise for reading in the fusiform gyrus. Trends in Cognitive Sciences 7:293–299.

Mercure E, Kadosh KC, Johnson MH (2011) The N170 shows differential repetition effects for faces, objects, and orthographic stimuli. Frontiers in Human Neuroscience 5:6.

Morel S, Ponz A, Mercier M, Vuilleumier P, George N (2009) EEG-MEG evidence for early differential repetion effects for fearful, happy and neutral faces. Brain Research 1254, 84–98.

Newsome WT, Pare EB (1988) A selective impairment of motion perception following lesions of the middle temporal visual area (MT). The Jouranl of Neuroscience 8:2201–2211.

Nichols TE, Holmes AP (2002) Nonparametric permutation tests for functional neuroimaging: A primer with examples. Human Brain Mapping 15:1–25.

Panichello MF, Cheung OS, Bar M (2013) Predictive feedback and conscious visual experience. Frontiers in psychology, 3, 620.

Pazo-Alvarez P, Cadaveira F, Amenedo E (2003) MMN in the visual modality: A review. Biological Psychology 63:199–236.

Perry G, Singh KD (2014) Localizing evoked and induced responses to faces using magnetoencephalography. European Journal of Neuroscience 39:1517–1527.

Righart R, de Gelder B (2006) Context influences early perceptual analysis of faces—an electrophysiological study. Cerebral Cortex 16:1249–1257.

Rossion B, Jacques C (2008) Does physical interstimulus variance account for early electrophysiological face sensitive responses in the human brain? Ten lessons on the N170. NeuroImage 39:1959–1979.

Rossion B, Caharel S (2011) ERP evidence for the speed of face categorization in the human brain: Disentangling the contribution of low-level visual cues from face perception. Vision Research 51:1297–1311.

Segaert K, Weber K, de Lange FP, Petersson KM, Hagoort P (2013) The suppression of repetition enhancement: a review of fMRI studies. Neuropsychologia, 51(1), 59–66.

Senju A, Johnson MH, Csibra G (2006) The development and neural basis of referential gaze perception. Social Neuroscience 1:220–234.

Simpson M, Johnson SR, Prendergast G, Kokkinakis AV, Johnson E, Green G, Johnston P (2015) MEG adaptation resolves the spatiotemporal characteristics of face-sensitive brain responses. The Jouranl of Neuroscience 35:15088–15096.

Stefanics G, Astikainen P, Czigler I (2014) Visual mismatch negativity (vMMN): A prediction error signal in the visual modality. Frontiers in Human Neuroscience 8:1074.

Summerfield C, Trittschuh EH, Monti JM, Mesulam MM, & Egner T (2008). Neural repetition suppression reflects fulfilled perceptual expectations. Nature neuroscience, 11(9), 1004–1006.

Summerfield C, de Lange FP (2014) Expectation in perceptual decision making: neural and computational mechanisms. Nature Reviews Neuroscience 15, 745–756

Tanaka JW, Curran T (2001) A neural basis for expert object recognition. Psychological Science 12:43–47.

Tian T, Feng X, Feng C, Gu R, Luo Y-J (2015) When rapid adaptation paradigm is not too rapid: Evidence of face-sensitive N170 adaptation effects. Biological Psychology 109:53–60.

Tipples J, Johnston P, Mayes A (2013) Electrophysiological responses to violation of expectation from eye gaze and arrow cues. Social Cognitive and Affective Neuroscience 8:509–514.

Tottenham N, Tanaka JW, Leon AC, McCarry T, Nurse M, Hare TA, Marcus DJ, Westerlund A, Casey B, Nelson C (2009) The NimStim set of facial expressions: Judgments from untrained research participants. Psychiatric Research 168:242–249.

Van Veen BD, Van Drongelen W, Yuchtman M, Suzuki A (1997) Localization of brain electrical activity via linearly constrained minimum variance spatial filtering. Biomedical Engineering, IEEE Transactions on Bio-medical Engineering 44:867–880.

Zhao L, Li J (2006) Visual mismatch negativity elicited by facial expressions under non-attentional condition. Neuroscience Letters 410:126–131.

